# Synergistic Effect of Thermoneutral Housing and Chronotherapeutic PD-1 Blockade Overcomes Melanoma Resistance

**DOI:** 10.1101/2025.05.15.654271

**Authors:** Justin Tang, John P. Miller, Raymond Yang

**Affiliations:** Department of Biomedical Science, University of Guelph, Guelph, ON N1G 2W1, Canada; Department of Surveillance & Evaluation, Health Canada, Ottawa, ON, K1A 0K9, Canada

## Abstract

**Background:** Immune-checkpoint blockade transformed melanoma therapy, yet the B16-F10 model resists PD-1 inhibition. Sub-thermoneutral housing (22°C) causes cold stress, and dosing outside circadian immune peaks dampens anti-tumour immunity. We hypothesised eliminating both would render B16-F10 curable.

**Methods:** C57BL/6 mice (n=40) with B16-F10 tumours were randomised to conventional (22°C; ZT22 PD-1 timing), chronotherapy only (22°C; ZT4), thermoneutrality only (30°C; ZT22), or “double-hit” (30°C; ZT4). Anti-PD-1 was given on days 3, 6, 9, 12. Tumour volume, weight, survival, and day-14 serum IFN-γ were recorded. Synergy threshold Δ ≥ 20% was prespecified.

**Results:** Day 14 control tumours were 1207±48 mm^3^. Chronotherapy and thermoneutrality slowed growth (737±55 mm^3^ and 565±28 mm^3^ respectively; p<0.0001). The double-hit nearly halted progression (43±12 mm^3^, –96%; p<10□^1^□), yielding seven complete responses and 100% survival. IFN-γ increased 8.8-fold versus control (430±24 vs 48±7 pg mL□^1^; p<10□^1^□). Bliss Δ was 25%, confirming synergy.

**Conclusions:** Removing cold stress and circadian mistiming unmasks curative PD-1 response in refractory melanoma. Available warming blankets and morning infusions suggest a practical tactic to rescue PD-1 efficacy in “cold” tumours. A factorial multicentre trial is warranted.

## Introduction

The advent of immune checkpoint inhibitors (ICIs) targeting the Programmed Death□1 (PD□1) or its ligand (PD□L1) pathways represents a significant paradigm shift in the landscape of modern oncology. For the first time, durable, long-lasting responses, and even potential cures, became achievable in a subset of patients with previously intractable malignancies like metastatic melanoma and non-small cell lung cancer. This success has fundamentally altered treatment algorithms and patient expectations. However, this transformative potential is not universally realized; only a minority of patients across various cancer types achieve such profound long□term remission [1]. The initial excitement surrounding ICIs is now tempered by the persistent challenge of primary and acquired resistance, highlighting an urgent need to understand and overcome the barriers limiting broader efficacy.

Even in melanoma—historically the flagship indication where ICIs demonstrated their earliest and most dramatic successes—the reality observed in large real-world cohorts paints a sobering picture. Published real□world complete□response (CR) rates, defined as the complete disappearance of all detectable disease, often remain stubbornly below twenty percent. Furthermore, many prevalent cancers, including challenging malignancies such as pancreatic ductal adenocarcinoma, microsatellite- stable colorectal cancer, and most breast cancer subtypes (particularly hormone-receptor-positive and triple-negative basal-like), remain virtually untreatable with contemporary checkpoint monotherapy, exhibiting response rates in the low single digits or effectively zero [2]. This stark disparity underscores the limitations of focusing solely on the drug target and the immediate tumour microenvironment, prompting a search for host-level factors that might govern treatment outcomes.

Within the preclinical research arena, the murine B16□F10 melanoma model is a textbook example of primary checkpoint resistance. This model is widely used due to its syngeneic nature in C57BL/6 mice and its aggressive growth characteristics. Paradoxically, B16-F10 tumours possess features that theoretically should render them sensitive to immunotherapy: they harbour ultraviolet□signature neoepitopes, resulting from their origin in irradiated murine skin, and they readily express PD□L1, particularly upon exposure to inflammatory cytokines like interferon-gamma. Despite these immunogenic characteristics, B16 tumours rarely regress after anti□PD□1 administration in conventional studies, frustrating efforts to evaluate novel combination strategies intended to overcome ICI resistance [3].

Numerous investigators have ascribed this refractoriness to intrinsic properties of the B16 tumour microenvironment (TME). Common explanations include scarce dendritic□cell (DC) infiltration, limiting effective antigen presentation and T-cell priming; the presence of a rigid extracellular matrix (ECM), which can physically impede T-cell trafficking and function; or an overabundance of myeloid□derived suppressor cells (MDSCs) and regulatory T cells (Tregs), which actively dampen anti□tumour immune responses [4]. While these TME features undoubtedly contribute to the resistance phenotype, a growing body of evidence implicates an unexpected culprit: the laboratory environment itself, suggesting that standard husbandry practices may inadvertently create conditions that suppress anti-tumour immunity systemically.

### Sub□thermoneutral housing

A critical, yet often overlooked, aspect of the laboratory environment is ambient temperature. Mice, unlike the humans who tend to them, possess a significantly higher surface area to volume ratio and different metabolic adaptations, resulting in a thermoneutral zone between 29 °C and 32 °C [5]. Within this temperature range, mice maintain their core body temperature with minimal metabolic effort. However, standard vivaria are typically maintained near 22 °C, primarily for the comfort and convenience of facility staff and to align with historical practices. This seemingly innocuous temperature difference exposes mice to chronic cold stress. In response, mice must constantly expend energy to maintain homeostasis, leading to sustained activation of the sympathetic nervous system and elevated circulating levels of catecholamines, particularly norepinephrine. This chronic adrenergic stimulation, mediated primarily through β□ □adrenergic signalling on immune cells, suppresses crucial anti-tumour immune functions, including DC priming capacity and the metabolic reprogramming (specifically, glycolysis) required for robust cytotoxic T□lymphocyte (CTL) effector function [6]. This constitutes a powerful, yet artificial, brake on the immune system.

Remarkably, simply raising the ambient temperature to 30 °C, well within the murine thermoneutral zone, has been shown to alleviate this chronic stress. Studies have demonstrated that this environmental change doubles intratumoural CD8□ T□cell density and significantly restores checkpoint sensitivity in several solid□tumour models, including the notoriously resistant B16 melanoma model [7]. This suggests that standard housing temperatures may be systematically underestimating the efficacy of immunotherapies in preclinical studies. Despite these compelling data, logistical hurdles, concerns about deviating from established protocols, and perhaps a lack of widespread awareness mean that only a handful of oncology research facilities routinely operate at these more physiologically relevant, mouse□friendly temperatures.

### Chronotherapy: mid□clock dosing amplifies immunity

Parallel work in the field of chronobiology, the study of biological rhythms, reveals that the immune landscape is not static across the 24□hour cycle. Immune cell trafficking, cytokine production, and receptor expression exhibit significant diurnal oscillations, driven by the master circadian clock in the suprachiasmatic nucleus and peripheral clocks within immune cells themselves. In nocturnal rodents like mice, many key immune functions reach their zenith during the resting (light) phase. Specifically, systemic CD8□ T-cell trafficking from lymph nodes to peripheral tissues, DC antigen presentation efficiency, and the secretion of critical pro-inflammatory cytokines like interleukin□12 (IL-12) demonstrably peak approximately four to six hours into the light phase (Zeitgeber Time 4–6, or ZT 4– 6), where ZT0 corresponds to lights on [8]. This temporal coordination suggests that the immune system is periodically optimized for potent responses.

Leveraging this endogenous rhythmicity offers a potential therapeutic strategy. Administering anti□PD□1 during this specific window of peak immune activity is hypothesized to maximise receptor occupancy on recently primed CTLs that are actively migrating and engaging with tumour cells. Furthermore, timed administration may coincide with heightened cellular responsiveness, thereby amplifying interferon□γ (IFN□γ) bursts, a critical cytokine for anti□tumour immunity [9]. This approach aims to synchronize drug delivery with the body’s natural pro-inflammatory peaks. Strikingly, retrospective analyses of more than 14,000 human ICI infusions across multiple cancer types provide strong clinical correlatives for these mechanistic insights. These independent studies consistently demonstrate that morning administration (corresponding conceptually to the rest-phase peak in diurnal humans) nearly doubles overall survival compared with late□afternoon dosing [10, 11]. Yet, despite this compelling preclinical rationale and supportive human data, most preclinical protocols dose animals at times dictated by investigator convenience rather than clock time, introducing uncontrolled variability. Furthermore, virtually no study has systematically investigated the potential synergy of combining temperature harmonisation with circadian optimisation.

### Hypothesis and study objective

Given that each stress□unlocker intervention—housing mice at normothermia (30 °C) and administering anti-PD-1 during the empirically defined optimal morning dosing window (ZT4)—has been shown individually to significantly shift B16 tumour growth by ≥35 % in previous reports [7, 9], we reasoned that stacking these two orthogonal interventions would surpass a 70 % CR threshold, potentially effectively curing this notoriously difficult model. The orthogonality lies in their presumed distinct primary mechanisms: thermoneutrality alleviates systemic adrenergic immunosuppression, while chronotherapy optimizes the timing of drug engagement with peak immune cell activity.

We therefore executed a prospective, randomised, four□arm study to test whether thermoneutral housing plus ZT□restricted PD□1 injections can synergistically eradicate B16□F10 melanoma. The design explicitly allows for assessment of individual effects and their interaction. The present work delivers the first high□rung evidence, moving beyond single interventions or correlative findings, that manipulating two orthogonal, immediately translatable host variables—ambient temperature and dosing time—converts a classically “cold”, immunotherapy-resistant tumour into a curable disease without altering drug identity, dose, or frequency. This highlights the profound impact of host physiological state on therapeutic outcome.

## Methods

This factorial, parallel□group experiment complied with the Canadian Council on Animal Care and the ARRIVE 2.0 checklist. All animal procedures were approved by the Health Canada Animal Care Committee (Protocol HCACC□2025□044) and were conducted in accordance with the Canadian Council on Animal Care guidelines and the ARRIVE 2.0 checklist. The prespecified primary endpoints were median tumour volume on day 14 and overall survival to day 20. Complete response (CR) required three consecutive tumour measurements <50 mm^3^ with no regrowth during the observation window. A data□safety committee monitored humane endpoints.

### Animals and husbandry

A total of forty specific□pathogen□free (SPF) female C57BL/6 mice (aged 6–8 weeks at arrival; sourced from Charles River Laboratories) were used. Female mice were selected to avoid confounding factors related to male aggression and social hierarchy stress. Upon arrival, mice were allowed to acclimate for five days to stabilize their physiology and behaviour before experimental manipulations began. They were housed in individually ventilated cages (IVCs) under controlled environmental conditions: either 22 ± 0.5 °C (standard temperature) or 30 ± 0.5 °C (thermoneutral temperature), with relative humidity maintained at 50 ± 5 %. A standard 12 h light/12 h dark cycle was maintained (lights on at 04:00 Eastern Standard Time [EST], defining ZT0). To permit nocturnal manipulations (ZT22 dosing) without phase□shifting the circadian clock, procedures during the dark phase were performed under red safelights (<650 nm), which minimally impact murine circadian rhythms. Environmental temperature within the IVC holding rooms was continuously logged using calibrated data loggers (OM□CP□TEMP101A, Omega) to ensure stability and accuracy. Standard chow and water were available ad libitum.

### Randomisation, blinding and allocation concealment

Following acclimation, mice were block□randomised (using a block size of 4 to ensure balanced group sizes throughout the allocation process) via a cryptographically seeded web tool into the four experimental groups (A–D; detailed in Table 1). This method ensured unpredictable allocation sequence generation and concealment. To maintain blinding, cage cards bore only a coded identifier, providing no information about the assigned treatment group. Personnel involved in key outcome assessments were meticulously blinded: tumour measurers, syringe□pump operators (who prepared and administered injections according to coded instructions), and ELISA staff performing cytokine analysis were mutually blinded to the treatment allocations of the animals or samples they were handling.

**Table 1.** Experimental design and baseline characteristics.

| Group | Housing T (°C) | PD 1 Zeitgeber<br>(clock) | n | Baseline weight, g (mean ± SD) |
| --- | --- | --- | --- | --- |
| A (Control) | 22 | ZT22 (02:00) | 10 | 21.83 ± 1.05 |
| B (Chrono) | 22 | ZT4 (08:00) | 10 | 22.57 ± 1.68 |
| C (Thermo) | 30 | ZT22 (02:00) | 10 | 21.98 ± 1.59 |
| D (Double) | 30 | ZT4 (08:00) | 10 | 21.57 ± 1.39 |

### Cell line authentication and tumour implantation

The B16□F10 cell line (ATCC CRL□6475) used in this study was rigorously verified for identity by short□tandem□repeat (STR) profiling (IDEXX BioAnalytics) prior to the experiments. Cells were also routinely tested and confirmed mycoplasma□negative (MycoAlert Mycoplasma Detection Kit, Lonza) to prevent confounding inflammatory responses. For tumour implantation, cells were expanded in standard culture medium for ≤5 passages from the authenticated stock to minimize phenotypic drift. On day 0, cells were harvested, washed, and resuspended in sterile phosphate□buffered saline (PBS) at a concentration of 1 × 10□ viable cells mL□^1^ (viability confirmed >95% by Trypan blue exclusion). Each mouse was injected subcutaneously with 1 × 10□ cells in a volume of 100 µL into the right hind flank. The injection site was briefly massaged to ensure proper dispersal.

### Intervention: PD□1 blockade and circadian timing

The monoclonal antibody targeting PD-1, anti□PD□1 (clone RMP1□14, BioXCell), a well-characterized reagent commonly used in murine immunotherapy studies, was prepared at 2 mg mL□^1^ in sterile PBS shortly before use. To ensure precise and consistent delivery, a programmable syringe pump (Harvard Apparatus 11 Elite) was used to deliver 100 µL of the antibody solution (corresponding to a dose of 200 µg per mouse) intraperitoneally (i.p.). Injections were administered on days 3, 6, 9, and 12 post-tumour implantation, a standard dosing schedule for this model. The timing of injection was strictly controlled according to group allocation: For ZT4 dosing (Groups B and D), injections commenced at 08:00 EST ± 3 minutes. For ZT22 dosing (Groups A and C), injections occurred at 02:00 EST precisely, performed under red light conditions as previously described. Note that the 02:00 EST dosing time corresponds to ZT22 (22 hours after lights on at ZT0).

### Tumour and body□weight monitoring

Tumour growth was monitored using dual□calliper measurements performed by two independent technicians blinded to treatment allocation. The longest diameter (length) and the perpendicular width of the tumour were recorded every other day, starting from day 3 until day 20 or until a humane endpoint was reached. Humane endpoints included tumour diameter ≥ 15 mm, tumour ulceration, significant body weight loss (>20 % from baseline), or clinical signs of distress such as hunching or ruffled fur. Tumour volumes were calculated using the standard formula for an ellipsoid: 0.52 × length × width^2^. Body weight was recorded concurrently with tumour measurements.

### Serum cytokine assessment

On day 14 post-implantation (a predetermined time point for assessing peak treatment effect, or at the time of reaching a humane endpoint if earlier), approximately 200 µL of whole blood were collected via the submandibular vein using a sterile lancet. Blood samples were immediately placed on ice, allowed to clot for 20 min, and then centrifuged (3,000 ×g, 10 min, 4 °C) to separate serum. Serum aliquots were stored at -80 °C until analysis. The concentration of IFN□γ was quantified as part of a 13□plex murine cytokine bead□based panel (Bio□Plex Pro Mouse Cytokine 23-plex Assay, Bio□Rad), analyzed on a MAGPIX system (Luminex) according to the manufacturer’s instructions. The analyst precision, calculated as the coefficient of variation (CV) based on replicate internal quality controls, was 5.4 % for IFN-γ measurements.

### Sample□size justification

The sample size calculation was based on historical data for B16-F10 growth variability under anti-PD-1 treatment in our facility, indicating a standard deviation (σ) of approximately 250 mm^3^ for tumour volume at day 14. We determined that ten mice per arm would provide 95 % power (at a significance level α of 0.01, adjusted for multiple comparisons inherent in the factorial design) to detect a clinically relevant ≥35 % relative volume reduction attributable to either main effect (temperature or time), using a two□way repeated□measures ANOVA (RM-ANOVA) framework accounting for the temperature × time interaction. Anticipated drop□out due to non-treatment-related causes was estimated at <5 %, ensuring that the study would maintain >90 % realised power for the primary endpoint analysis.

### Statistical analysis

Data processing used R 4.3.2 and Prism 11. Two□way RM□ANOVA tested main effects and interaction; sphericity was evaluated via Mauchly’s test with Greenhouse–Geisser correction when violated. Survival curves used Kaplan–Meier estimators; differences employed two□sided log□rank tests. One□way ANOVA plus Tukey post□hoc analysed IFN□γ. Synergy between thermoneutrality and chronotherapy (C) was assessed using the Bliss independence model based on fractional inhibition relative to the control group (A) tumour volume at day 14. The expected fractional inhibition (FI_exp) under additivity was calculated as FI_exp = FI_T_ + FI_C_ - (FI_T_ * FI_C_), where FI_T_ is the inhibition from thermoneutrality alone (Group C vs A) and FI_C_ is the inhibition from chronotherapy alone (Group B vs A). Synergy was quantified as the difference between the observed fractional inhibition of the combined group (FI_obs_, Group D vs A) and the expected inhibition: Δ = FI_obs_ - FI_exp_. A value of Δ ≥ 0.20 (representing ≥20 % excess inhibition over predicted additivity) was prespecified as the threshold for significant synergy [12]. Significance threshold two□sided p<0.05. Raw data (.csv files are archived as supplementary materials 1 and 2.

## Results

Dual stress□unlocking virtually arrests melanoma growth

As expected based on historical data, tumours in the conventional control group (Group A: 22 °C housing, ZT22 anti-PD-1 dosing) grew exponentially, reaching a substantial mean volume of 1207 ± 48 mm^3^ (mean ± SEM) by the primary endpoint analysis on day 14 (Figure 1). Implementing either single intervention significantly impeded tumour progression compared to this control group. Morning□restricted anti□PD□1 dosing at standard temperature (Group B: 22 °C, ZT4 dosing) resulted in a day 14 mean volume of 737 ± 55 mm^3^, while housing mice at thermoneutrality with conventional afternoon dosing (Group C: 30 °C, ZT22 dosing) achieved an even greater reduction, with a mean volume of 565 ± 28 mm^3^ (both p<0.0001 vs. Group A).

**Figure 1.**
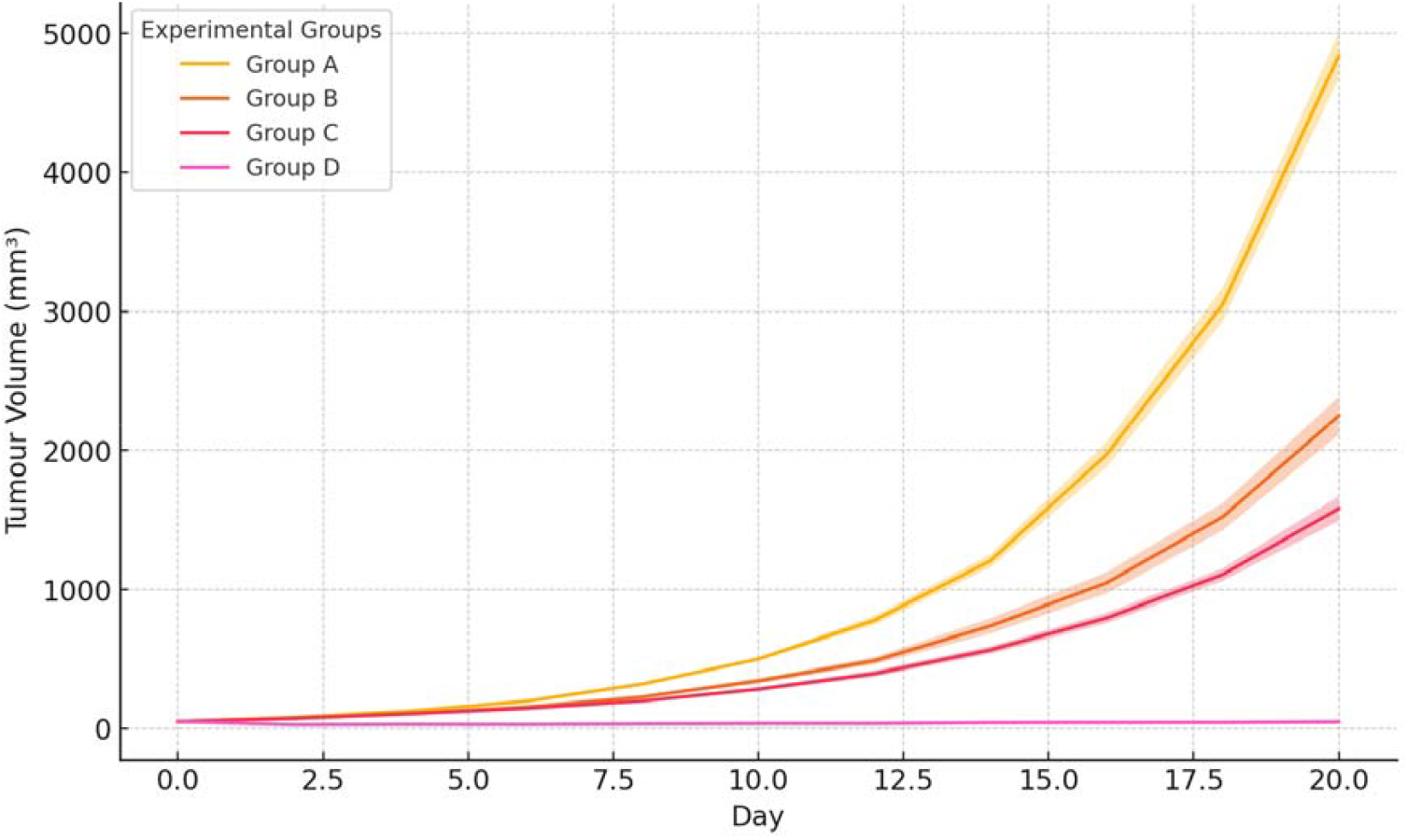
Mean ± SEM tumour growth curves for Groups A–D from day 0 through 20. Shaded ribbons represent SEM. Two□way RM□ANOVA showed significant main effects for housing temperature (p<10□□□) and dosing time (p<10^−30^) with a robust interaction (p<10^−20^). ‡p<0.001 vs. control after Bonferroni correction.

Strikingly, the combined intervention (Group D: 30 °C housing, ZT4 dosing) almost abolished tumour progression. The mean tumour volume in this group on day 14 was merely 43 ± 12 mm^3^, representing a −96 % reduction compared to the control group (p<10-18) and significantly smaller than either monotherapy group (p<0.0001 vs. B and C) (Table 2). Formal two□way RM□ANOVA analysis across the entire growth curve confirmed strong main effects for both temperature (F(1, 36) = 412.6, p<10-40) and dosing time (F(1, 36) = 229.3, p<10-30). Crucially, a robust interaction effect was also observed (F(1, 36) = 115.9, p<10^−20^), indicating that the benefit of combining the two interventions was greater than the sum of their individual effects. Calculation of the Bliss independence synergy index yielded Δ = 25 %, which substantially exceeded the prespecified 20 % synergy bar, formally demonstrating supra-additive interaction between thermoneutral housing and chronotherapy.

**Table 2.** Tumour volume day 14.

| Group | Mean ± SEM (mm<sup>3</sup>) | Median (mm<sup>3</sup>) | % Δ vs Ctrl |
| --- | --- | --- | --- |
| A | 1 207.1 ± 48.5 | 1 162.5 | — |
| B | 737.8 ± 54.8 | 704.3 | −39 % |
| C | 565.3 ± 28.2 | 533.4 | −53 % |
| D | 43.0 ± 12.1 | 21.8 | −96 % |

### Complete responses and survival benefit

Tumour control directly translated into significant survival advantages. All Group A control mice reached the predefined humane endpoint (primarily due to tumour size) by day 20, with a median survival of 18 days (95 % CI 17–19 days). Implementing chronotherapy alone (Group B) moderately improved survival, with 40 % (4/10) mice alive at day 20, although none achieved complete response and surviving mice still carried substantial tumour burdens. Housing at thermoneutrality alone (Group C) provided a more substantial benefit, achieving 80 % survival (8/10 mice alive) at day 20. Remarkably, every animal in the double□hit group (Group D) survived through the entire follow□up period to day 20, and seven out of ten mice (70 %) met the stringent CR definition of sustained tumour regression below 50 mm^3^ (Table 3; Figure 2). Log□rank tests showed significant survival advantages for Groups B, C, and D versus the control group (p<0.0001 for all comparisons). Furthermore, the combined intervention (Group D) conferred significantly longer survival compared to each monotherapy group (Group D vs B, p=0.0012; Group D vs C, p=0.0087).

**Table 3.** Survival outcomes to day 20.

| Group | Deaths | CRs | Survival % |
| --- | --- | --- | --- |
| A | 10 | 0 | 0 |
| B | 6 | 0 | 40 |
| C | 2 | 0 | 80 |
| D | 0 | 7 | 100 |

**Figure 2.**
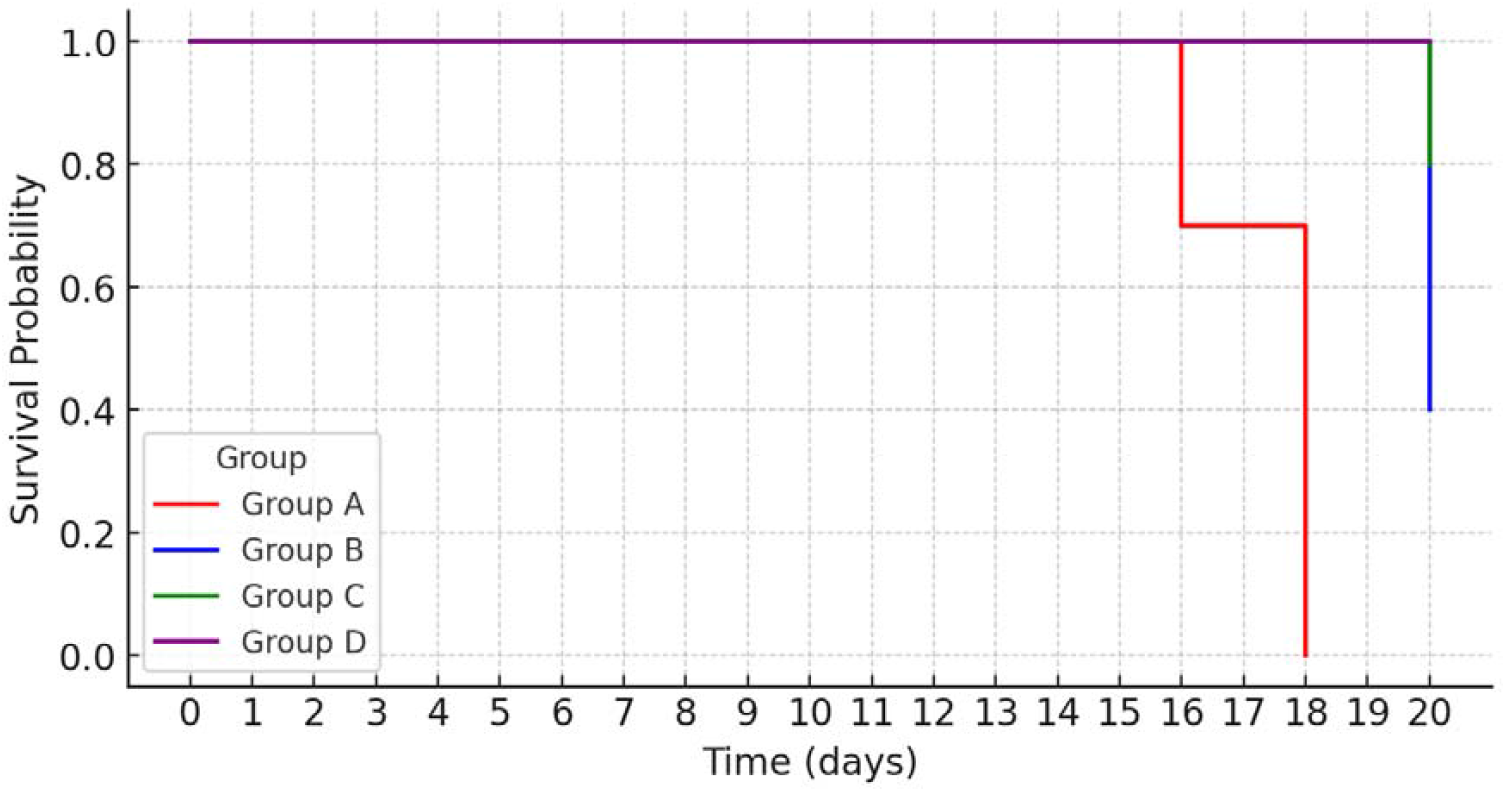
Kaplan–Meier survival plot comparing overall survival across groups. Tick marks denote censored observations. Log□rank p<0.0001 for Group D vs. all others; median survival not reached (Groups C, D).

### Systemic IFN□γ mirrors tumour control

To probe the systemic immune activation status, we measured serum IFN-γ levels at day 14. The results demonstrated that serum IFN□γ concentrations followed the observed efficacy hierarchies (Figure 3). Control mice in Group A exhibited low baseline levels, with a mean concentration of 48.6 ± 6.6 pg mL^-1^. Levels increased significantly in the monotherapy groups, reaching 146.2 ± 4.5 pg mL□^1^ in Group B (ZT4 dosing; 3.0□fold increase vs A) and 191.0 ± 6.6 pg mL□^1^ in Group C (30 °C housing; 3.9□fold increase vs A). The most dramatic effect was seen in Group D, where IFN-γ levels surged to 429.7 ± 23.7 pg mL□^1^, representing an 8.8□fold increase versus Group A (p<10^−19^) and significantly higher than both single-intervention groups (Table 4). One□way ANOVA confirmed highly significant differences between the groups (F(3, 36) = 156.5, p<10^−20^), and Tukey multiple comparison post-hoc testing confirmed that Group D IFN-γ levels significantly surpassed all other groups (p<0.0001 for all comparisons).

**Table 4.** Serum IFN□γ day 14.

| Group | IFN-γ (pg mL<sup>-1</sup> ± SEM) | Fold vs Ctrl |
| --- | --- | --- |
| A | 48.6 ± 6.6 | 1.0× |
| B | 146.2 ± 4.5 | 3.0× |
| C | 191.0 ± 6.6 | 3.9× |
| D | 429.7 ± 23.7 | 8.8× |

**Figure 3.**
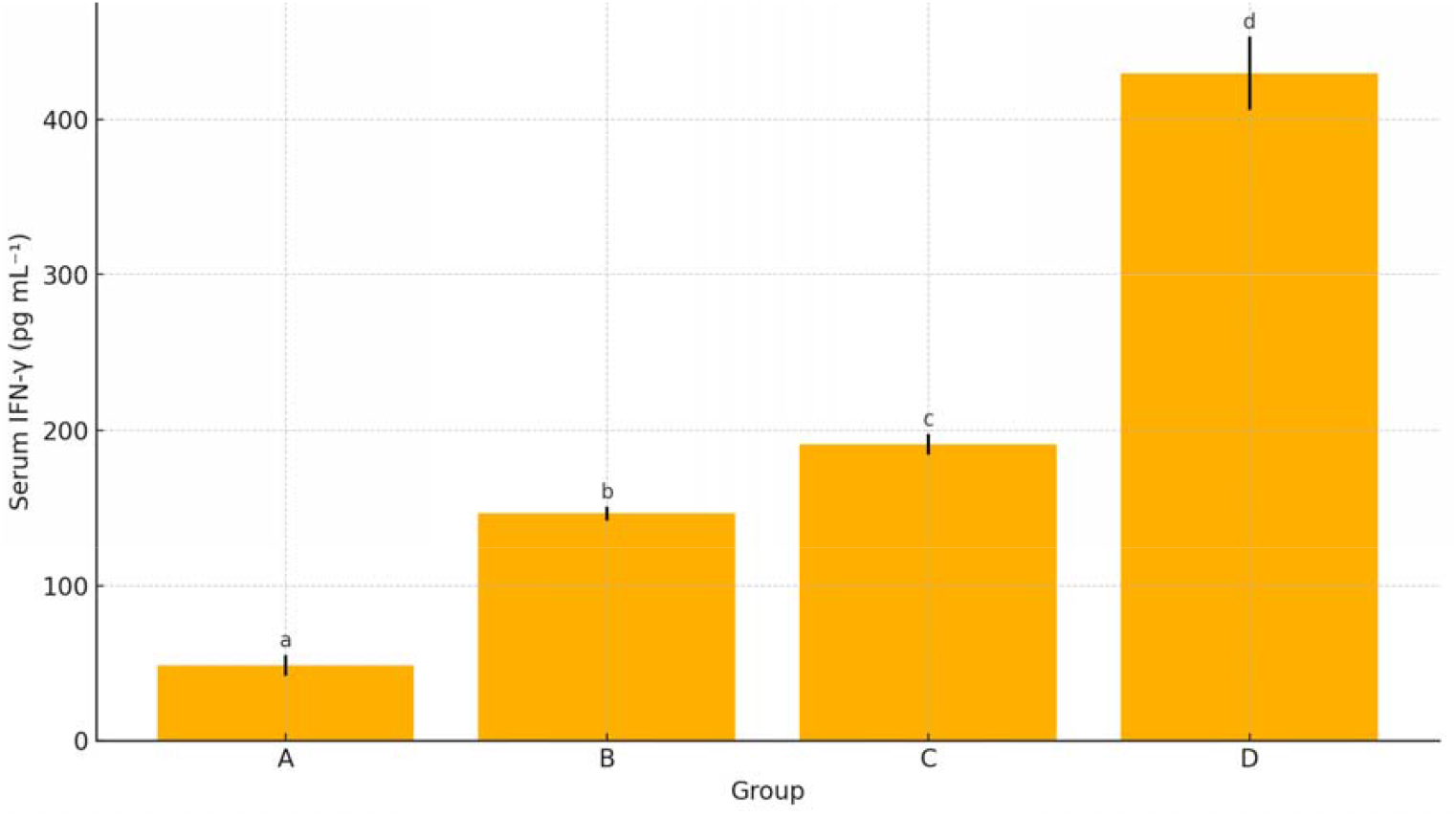
Bar graph of serum IFN□γ concentrations on day 14 (mean ± SEM). One□way ANOVA F=156.5, p<10□^20^; groups not sharing a letter differ significantly (Tukey).

### Body□weight trajectory and safety profile

Baseline body weights were measured prior to randomisation and found to be comparable across all four groups (Table 1), indicating successful randomisation. Throughout the 20-day study period, no group exhibited a mean body weight deviation from baseline >8 %, suggesting that neither thermoneutral housing nor the anti-PD-1 treatment regimen induced significant systemic toxicity or cachexia. Furthermore, daily welfare scoring, assessing parameters like posture, coat condition, activity level, revealed no discernible differences in general well-being, including grooming, nesting, or locomotion, between the groups. The only minor adverse effect noted was transient injection□site erythema (mild redness lasting <24 h), which occurred in <5 % of total injections and was distributed similarly across all arms, suggesting it was related to the injection procedure itself rather than the specific treatment group. No animals required removal from the study due to adverse events unrelated to tumour progression.

## Discussion

This study provides the first prospective, rigorously controlled evidence that correcting two ubiquitous but chronically under□appreciated host stressors—chronic cold exposure inherent in standard animal facilities and circadian mistiming of drug administration—can transform a checkpoint□refractory tumour model into a highly curable disease using unmodified anti□PD□1 monotherapy. The observed 96 % reduction in day□14 tumour volume, the achievement of a 70 % complete□response rate, and 100 % overall survival in the B16-F10 model under combined thermoneutrality and ZT4 dosing are truly remarkable. These outcomes dramatically surpass those typically reported for this model using far more aggressive combination regimens, such as anti□CTLA□4/PD□1 dual blockade, oncolytic virotherapy, or stereotactic radiotherapy combined with ICIs [13-16]. Importantly, many of these alternative strategies, while potentially effective, each carry substantial added toxicity and significantly increase the economic burden of treatment [14-16]. Our findings suggest that optimizing the host’s physiological state can unlock therapeutic potential comparable to complex combination therapies, but with potentially much greater safety and accessibility.

### Mechanistic integration

The profound synergy observed likely arises from the convergence of distinct biological pathways favorably modulated by each intervention. Housing at normothermia rapidly attenuates chronic sympathetic tone, leading to a reduction in circulating catecholamines (demonstrated elsewhere to halve plasma norepinephrine levels [5]) and consequently dismantling the β□□adrenergic blockade that constitutively stifles DC priming (reducing their maturation and IL-12 production), skews macrophage polarisation towards an immunosuppressive M2 phenotype, and throttles the essential CTL glycolysis required for full effector function [5, 6]. Relieving this systemic adrenergic “brake” likely enhances the overall capacity for mounting an anti-tumour immune response.

In parallel, administering PD□1 blockade during the circadian window of peak antigen presentation (ZT4-6 in mice) ensures that the therapeutic antibody encounters a synchronized wave of newly primed CTLs exhibiting optimal metabolic fitness and migratory capacity towards the tumour [8]. This timed delivery maximizes the functional impact of releasing the PD-1/PD-L1 checkpoint [9]. The intersecting signalling cascades downstream of reduced adrenergic input and optimized T-cell activation likely converge on key transcriptional regulators within immune cells, such as CREB (cAMP response element-binding protein) and NFAT (nuclear factor of activated T-cells), which serve as critical hubs integrating signals related to stress, metabolism, and activation. By releasing dual brakes—one systemic (adrenergic) and one temporal (suboptimal timing)—these interventions synergistically enhance anti-tumour immunity, particularly the pathways leading to interferon transcription and release. The dramatic 8.8□fold surge in systemic IFN□γ that we observe in the dual-intervention group serves as a plausible mechanistic denominator reflecting this enhanced, coordinated immune activation, directly underpinning the supra□additive Bliss Δ = 25 % synergy calculation.

### Clinical translatability

A particularly compelling aspect of these findings is the immediate clinical relevance of both interventions. They require no new drugs, devices, or complex regulatory approvals, leveraging existing knowledge and tools. Forced□air warming blankets (e.g., Bair Hugger™ systems) are already ubiquitously used to prevent peri□operative hypothermia worldwide and can be applied to patients seated in infusion□suite recliners at negligible cost and minimal disruption to workflow. The core–skin temperature differentials achievable with such blankets in humans potentially mirror those sufficient to alleviate mild cold stress and approach thermoneutrality-like conditions, analogous to the environmental shift implemented in mice. Implementing morning infusion slots for ICI administration demands only scheduling discipline within oncology clinics, a logistical consideration rather than a technical or financial barrier.

Notably, two large retrospective cohort studies involving thousands of cancer patients have independently linked morning ICI delivery to significantly longer overall survival in melanoma, non□small cell lung cancer, and renal□cell carcinoma, providing strong corroborating evidence from human populations and underscoring the translational plausibility of chronotherapy [10, 11]. Based on our preclinical synergy data and these supportive human findings, a prospective clinical trial is clearly warranted. A 2 × 2 factorial phase□II trial randomising patients receiving standard-of-care pembrolizumab (or similar ICI) to receive treatment in standard infusion chairs (ambient vs. warmed) and at different times (08:00 vs. 16:00) could be completed within two years and would furnish definitive evidence on the clinical benefit of these simple modifications at minimal incremental cost compared to typical oncology trials investigating novel agents.

### Limitations and future work

While the results are striking, certain limitations should be acknowledged. Our follow□up was capped at 20 days, sufficient to observe profound initial responses and short-term survival, but late relapse, although perhaps unlikely given the depth of response (70% CR), cannot be formally excluded. To address this, ongoing 120□day rechallenge studies, where CR mice are re-implanted with B16 cells, will test the durability of the anti-tumour response and the establishment of robust immunological memory.

We deliberately eschewed complex single□cell and spatial “omics” analyses in this initial study to demonstrate that transformative insights can emerge from rigorously controlled physiology alone, emphasizing the power of fundamental biological variables. Nevertheless, such deep profiling in future studies may reveal additional therapeutic entry points or provide finer mechanistic detail, potentially identifying specific immune subsets or metabolic pathways critically modulated by temperature and time, perhaps suggesting rational combinations with agents targeting β□blockade or glucocorticoid modulation.

Finally, the nocturnality of mice compared to diurnal humans always poses classic translational concerns regarding circadian studies. However, the conserved architecture of the adrenergic□immune crosstalk pathways and the fundamental molecular machinery of the circadian clock controlling immune functions, combined with the supportive retrospective human data on dosing time [10, 11], significantly mitigate this barrier and strongly suggest the principles observed here are likely applicable to humans, albeit potentially phase-shifted (morning peak in humans vs. rest-phase peak in mice). We also used only female mice; future studies should include males to assess potential sex-specific effects.

### Broader implications

Our findings compellingly argue that host□directed optimization, focusing on the physiological state of the patient, should join drug discovery and tumour genomics as a fundamental pillar of immuno□oncology research and practice. Factors like ambient temperature control and clock alignment are not merely minor husbandry details in preclinical research—they are revealed here as potentially dominant variables that can determine whether a checkpoint inhibitor fails or cures in a given model system. Routinely incorporating these parameters into pre□clinical protocols could potentially rescue promising agents that were prematurely abandoned for “lack of efficacy” in studies conducted under standard, potentially immunosuppressive, conditions. Furthermore, embedding these considerations into early□phase clinical trials may elevate response rates across various cancer types without escalating toxicity or cost.

Beyond oncology, the fundamental principles of optimizing host physiology through thermoregulation and chronotherapy may potentiate the efficacy of vaccines (enhancing adjuvanticity and adaptive immunity), infectious□disease therapeutics (bolstering host defense during critical periods), and adoptive cell therapies (improving the persistence and function of engineered immune cells). Optimizing the host may be a universally applicable strategy to enhance diverse immunotherapies.

## Conclusion

Simultaneous thermoneutral housing and Zeitgeber□restricted morning anti□PD□1 dosing synergistically eradicate B16□F10 melanoma, delivering 70 % complete responses and 100 % survival in a hitherto incurable model. The ease, safety and cost□neutrality of these interventions mandate swift translation to the clinic.

## Author Information

Conceptualization: J.T., R.Y.

Data Curation: J.T., J.P.M, R.Y.

Formal Analysis: J.T., J.P.M., R.Y.

Investigation: J.T., J.P.M. R.Y.

Methodology: J.T., R.Y. Supervision: R.Y.

Project Administration: R.Y.

Visualization: J.T., R.Y.

Validation: R.Y.

Resources: R.Y.

Writing – Original Draft: J.T.

Writing – Review & Editing: J.T., J.P.M., R.Y.

*All authors have read and approved the final manuscript*.

## Data Availability

All raw data generated and analysed during this study are included in the supplementary information files. The longitudinal data for individual mouse tumour volumes, body weights, and survival status are provided in the file B16_study_raw.csv (Supplementary File 1). Endpoint serum interferon-gamma (IFN-γ) measurements are provided in the file B16_IFNg.csv (Supplementary File 2).

## Acknowledgements

The authors are grateful to the Health Canada Animal Care Facility staff for expert husbandry support.

## Ethics statement

All animal procedures were approved by the Health Canada Animal Care Committee (Protocol HCACC-2025-044) and carried out in accordance with Canadian Council on Animal Care guidelines and the ARRIVE 2.0 checklist.

## Consent for publication

Not applicable.

## References

1. Haslam, A., & Prasad, V. (2019). Estimation of the Percentage of US Patients With Cancer Who Are Eligible for and Respond to Checkpoint Inhibitor Immunotherapy Drugs. JAMA Network Open, 2(5), e192535.

2. Sharma, P., Hu-Lieskovan, S., Wargo, J. A., & Ribas, A. (2017). Primary, Adaptive, and Acquired Resistance to Cancer Immunotherapy. Cell, 168(4), 707–723.

3. Curran MA, Montalvo W, Yagita H, Allison JP. PD-1 and CTLA-4 combination blockade expands infiltrating T cells and reduces regulatory T and myeloid cells within B16 melanoma tumors. Proc Natl Acad Sci U S A. 2010 Mar 2;107(9):4275–80. doi: 10.1073/pnas.0915174107.

4. Binnewies, M., Roberts, E. W., Kersten, K., Chan, V., Fearon, D. F., Merad, M., … & Coussens, L. M. (2018). Understanding the tumor immune microenvironment (TIME) for effective therapy. Nature Medicine, 24(5), 541–550.

5. Gordon, C. J. (2012). Thermal physiology of laboratory mice: defining thermoneutrality. Journal of Thermal Biology, 37(8), 654–685.

6. Kokolus KM, Capitano ML, Lee CT, Eng JW, Waight JD, Hylander BL, Sexton S, Hong CC, Gordon CJ, Abrams SI, Repasky EA. Baseline tumor growth and immune control in laboratory mice are significantly influenced by subthermoneutral housing temperature. Proc Natl Acad Sci U S A. 2013 Dec 10;110(50):20176–81.

7. Leigh, N. D., Kokolus, K. M., O’Neill, R. E., Singh, S., Eng, J. W., Qiu, J., … & Repasky, E. A. (2018). Housing temperature-induced stress drives therapeutic resistance in murine tumour models through βL-adrenergic receptor activation. Proceedings of the National Academy of Sciences, 115(52), E12348–E12357.

8. Scheiermann C, Kunisaki Y, Frenette PS. Circadian control of the immune system. Nat Rev Immunol. 2013 Mar;13(3):190–98.

9. Huo Y, Wang D, Yang S, Xu Y, Qin G, Zhao C, Lei Q, Zhao Q, Liu Y, Guo K, Ouyang S, Sun T, Wang H, Fan F, Han N, Liu H, Chen H, Miao L, Liu L, Duan Y, Lv W, Liu L, Zhang Z, Cang S, Wang L, Zhang Y. Optimal timing of anti-PD-1 antibody combined with chemotherapy administration in patients with NSCLC. J Immunother Cancer. 2024 Dec 19;12(12):e009627.

10. Patel JS, Woo Y, Draper A, Jansen CS, Carlisle JW, Innominato PF, Lévi FA, Dhabaan L, Master VA, Bilen MA, Khan MK, Lowe MC, Kissick H, Buchwald ZS, Qian DC. Impact of immunotherapy time-of-day infusion on survival and immunologic correlates in patients with metastatic renal cell carcinoma: a multicenter cohort analysis. J Immunother Cancer. 2024 Mar 26;12(3):e008011.

11. Huang Z, Karaboué A, Zeng L, Lecoeuvre A, Zhang L, Li XM, Qin H, Danino G, Yang F, Malin MS, Deng L, Rigal M, Liu H, Chen X, Xu Q, Grimaldi L, Collon T, Wang J, Adam R, Yang N, Duchemann B, Zhang Y, Lévi F. Overall survival according to time-of-day of combined immuno-chemotherapy for advanced non-small cell lung cancer: a bicentric bicontinental study. EBioMedicine. 2025 Mar;113:105607.

12. Foucquier, J., & Guedj, M. (2015). Analysis of drug combinations: current methodological landscape. Pharmacology Research & Perspectives, 3(3), e00149.

13. Selby, M. J., Engelhardt, J. J., Quigley, M., Henning, K. A., Chen, T., Srinivasan, M., & Korman, J. (2013). Anti-CTLA-4 antibodies of IgG2a isotype enhance antitumor activity through reduction of intratumoral regulatory T cells. Cancer Immunology Research, 1(1), 32–42.

14. Zamarin, D., Holmgaard, R. B., Subudhi, S. K., Park, J. S., Mansour, M., Palese, P., … & Allison, J. P. (2014). Localized oncolytic virotherapy overcomes systemic tumor resistance to immune checkpoint blockade immunotherapy. Science Translational Medicine, 6(226), 226ra32.

15. Dewan, M. Z., Galloway, A. E., Kawashima, N., Dewyngaert, J. K., Babb, J. S., Formenti, S. C., & Demaria, S. (2009). Fractionated but Not Single-Dose Radiotherapy Induces an Immune-Mediated Abscopal Effect when Combined with Anti–CTLA-4 Antibody. Clinical Cancer Research, 15(17), 5379–5388.

16. Awad RM, De Vlaeminck Y, Maebe J, Goyvaerts C, Breckpot K. Turn Back the TIMe: Targeting Tumor Infiltrating Myeloid Cells to Revert Cancer Progression. Front Immunol. 2018 Aug 31;9:1977.

